# Adult Neuronal Expression of the Alzheimer Risk Protein CD2AP is Enriched in NGF-Responsive Basal Forebrain Neurons, where it Collocates with Rab-5 Endosomes

**DOI:** 10.1101/2024.07.24.604961

**Authors:** Lindsey Avery Fitzsimons, Mohammad Atif-Sheikh, Madison Mueth, Makaela Rice, Gareth Howell, Benjamin J. Harrison

## Abstract

Genome-wide association studies (GWAS) with multiple human populations have uncovered single nucleotide variants of the *CD2AP* gene locus that are associated with Alzheimer’s Disease (AD) risk. However, the role of CD2AP in AD pathogenesis remains unknown. In adult PNS neurons, previous work demonstrated that CD2AP functions as a docking-scaffold/adaptor protein coordinator of nerve growth factor (NGF) trophic signaling and RAB5-mediated endocytosis. In the adult CNS, whereas CD2AP is robustly expressed in non-neuronal cells and neurovasculature, neuronal expression is restricted and poorly characterized. In this study using publicly available single cell/nucleus RNA sequencing, we observed that CD2AP mRNA is enriched in a population of TrkA-expressing cholinergic neurons of the adult mouse basal forebrain. Immunohistology using brain tissue from adult choline acetyltransferase (ChAT) GFP reporter mice, confirmed enrichment of CD2AP protein in cholinergic projections, and in neuron soma in the diagonal band of Broca where it co-localized with RAB5. Together with previous studies from NGF-responsive PNS primary sensory neurons, these observations indicate that CD2AP may play a role in retrograde trophic signaling in NGF-responsive CNS cholinergic neurons. In addition, we observed that CD2AP expression in these soma was increased in aged mice (18-month-old), concomitant with a reduction of co-localization with RAB5, suggesting a potential role for CD2AP in aging-associated changes in retrograde trophic signaling in these neurons. Basal forebrain cholinergic neurons project to the hippocampus and cortex, are required for learning and memory, are critical for brain health during aging, and disruption of RAB5 mediated endocytosis in these neurons is central to the pathogenesis of Alzheimer’s disease (AD). Future studies are therefore warranted to determine if CD2AP risk variants impact endocytosis and trophic signaling in cholinergic neurons in healthy aging and/or AD.

## Introduction

CD2-associated protein (CD2AP) is a docking-scaffold/adaptor protein with multiple binding interfaces for coordination of protein-protein interactions. It was first characterized as an SH-3- containing protein that binds to CD-2 via the cytoplasmic domain^1–3^. CD2AP serves as a hub for signaling pathway components that converge on this scaffold. When the required combination of interactions are formed, the corresponding downstream signaling cascade is activated, and for example, the presence or absence of CD2AP-binding interactions differentially modulates the amplitude of MAPK-ERK or PI3K signal transduction^4^. Known upstream binding-partners include Tropomyosin receptor kinase-A (TrkA) and/or rearranged during transfection (RET); activated by nerve growth factor (NGF) and glial cell line-derived neurotrophic factor (GDNF) family ligands (GFLs)^5,6^. Known downstream effector binding-partners include the PI3K regulatory subunit p85^5^, p21ras^7^ and Akt^8^.

Neurons are highly compartmentalized, and rely on retrograde trophic signaling though endosomal transport of internalized ligand-receptor complexes towards downstream effectors located in the cell body^9–11^. Variants in genes associated with endocytosis may be protective, e.g., associated with resilience to Alzheimer’s Disease (AD)^12^. Conversely, pathogenic variants have been elucidated by genome-wide association studies (GWAS), as factors associated with increased susceptibility to AD^12,13^. Specifically, the *Cd2ap* gene is an AD-predisposition locus, substantiated by multiple GWAS with multiple populations (African American^14,15^, Chinese^16^, Northern and Western European ancestry (CEU)^17^, UK Biobank^18,19^) that support the association of *Cd2ap* variants with small but significant increases in incidence^20^, progression^20,21^, and severity^16,21^ of Alzheimer’s Disease (AD).

CD2AP is expressed by multiple cell types in multiple tissues^22^, and may play a role in several mechanisms responsible for the pathogenesis and/or progression of AD. For example, dysfunction of brain vasculature is a pathophysiological mechanism of blood brain barrier deficits^23^ relevant to multiple dementias, including AD, (where CD2AP highly expressed)^24–26^. Outside of the vasculature, CD2AP is expressed by migratory dendritic cell populations^22,27^, renal podocytes^22,28^, and sensory neurons of the peripheral nervous system^5^. Thus, the clinical importance of *Cd2ap* is wide-ranging. However, the mechanistic underpinning of *Cd2ap* variant-associated risk involving the dynamic interplay between/among cell and tissue type systems, remains poorly characterized.

Phosphorylated Tau protein, forming neurofibrillary tangles, impedes conductance, retrograde signaling and therefore function of neurons, resulting in the neurotoxicity and degradation seen in healthy aging as well as in neurodegenerative disease like AD^29,30^. A genetic association between CD2AP, tau-accumulation and subsequent neurotoxicity^31^ have been observed using drosophila homolog *cindr in a* loss of function model of AD^31,32^. CD2AP scaffolding interactions regulate Rab5- mediated endocytosis^2,17,33–38^. Therefore, reduced CD2AP function may lead to impaired clearance and subsequent accumulation of amyloid-beta (Aß) and tau aggregates^33^.

Histological assessment of CD2AP expression in nervous tissues reveals strong expression in neurovascular. However, expression of CD2AP in adult neurons remains poorly characterized. CD2AP has been shown to be expressed in embryonic dorsal root ganglia (DRG) in E11 mouse embryos^39^, in adult mouse brain cerebella neurons and adult DRG^5^. In neuron-like PC12 cancer cell lines and adult mouse DRG, NGF stimulates CD2AP binding to Neurotrophic Receptor Tyrosine Kinase 1 (Gene name *Ntrk1*, Protein name: Tropomyosin Receptor Kinase A (TrkA)) and the PI3- kinase effector (p85), to upregulate the PI3K/AKT pathway, and is a positive coordinator of NGF-stimulated axon growth and branching^5^. Experiments investigating the Ras and Rab Interactor 3 (RIN3), observed that CD2AP is present in cultured murine cholinergic neurons isolated from basal forebrain, and that endosomal trafficking involving CD2AP *in vitro*, is reliant upon RIN3, a guanine nucleotide exchange factor (GEF) in the Rab5 small GTPase family^40^. CD2AP has also been characterized in the nervous system of *Drosophila, via its* homologue, *cindr*^41^. *Cindr* expression can be found in adult Drosophila throughout the brain/nervous system, and is predominantly expressed within neuropil and the surrounding presynaptic terminals, where it is shown to function as a positive regulator and rate-limiting component of synaptic transmission, calcium release, and proteostasis^41^. Silencing of *cindr* (CD2AP) robustly enhanced tau-toxicity in Drosophila^32^, using a human mutant form of Tau shown to be associated with familial frontotemporal dementia ^42^.

Previously, we observed that CD2AP is expressed in primary sensory neurons, and demonstrated that this adaptor protein coordinates NGF-TrkA-PI3K trophic signaling and retrograde transport via Rab5-decorated endosomes^5^. In the current study, we observed that in neurons of the central nervous system (CNS), CD2AP expression is highly enriched in a small population of cholinergic neurons in the basal forebrain and striatum. Cholinergic projections from the basal forebrain to the cortex constitute the primary source of cholinergic input to the forebrain^43^, and are also known to be the first to show signs of degeneration in AD^44^. Disruption of NGF-TrkA trophic signaling via interruption of retrograde Rab5a endosomes is a key mechanism of AD pathogenesis^35^. We therefore examine the localization of CD2AP with TrkA with Rab5a endosomes in cholinergic neurons and discuss a possible role for CD2AP in the trophic support of cholinergic neurons and AD pathophysiology.

## Materials & Methods Animals

Animal housing and procedures were approved by the Institutional Animal Care and Use Committee of the University of New England consistent with federal regulations and guidelines. All animals used for this study were ChAT^BAC^-eGFP mice (JAX Strain# 007902) purchased from The Jackson Laboratory (Bar Harbor, Maine). Age and sex of cohorts are specified in appropriate sections below.

### 2.2 Prediction of CD2AP mRNA Expression in Neuronal Populations

The DropViz online search tool was used to uncover putative populations of *Cd2ap*-expressing adult mouse neurons ^45^. The “Limit by Class” option was set to “Neuron” and the top-10 results were retrieved using following individual search terms: “*Cd2ap*”, “*Ntrk1*” and “*Chat*”. The top-10 CD2AP-expressing populations were selected.

### 2.3 Tissue Preparation & Frozen Tissue Sectioning

Six week-old, ChAT-GFP positive mice (N=2 males, 2 females), were deeply anaesthetized with 0.1ml intraperitoneal Euthasol, before intracardial perfusion with 20ml phosphate buffered saline (PBS) followed by 30ml cold 4% paraformaldehyde in PBS. Brains were immediately harvested, and post fixed in cold 4% paraformaldehyde in PBS for 4 hours. Following fixation, tissues were washed with PBS and incubated in 30% sucrose cryoprotectant for 48 hours. Tissues were then embedded in OCT frozen tissue embedding medium and stored at -80C prior to sectioning. Coronal sectioning of mouse brains embedded in OCT was performed using a Leica Cryostat 1900 (Leica Microsystems, USA). From each mouse, fifteen 12um thick, coronal brain sections were sliced 1.77mm to 0.13mm rostral to the bregma onto a glass slide (VWR®, Superfrost® Plus Micro Slide, Cat# 48311-703).

### 2.4 Tissue Processing and Immunofluorescent Staining

Sections were dried overnight, then briefly washed with PBS before incubation with blocking solution (4% normal donkey serum, 0.3% Triton X-100 PBS) for 30 mins at room temperature. Following blocking, slides were incubated with primary antibodies at the indicated dilutions (Table 1) over-night (18 hours) at room temperature. Slides were then washed 5 times for 5 minutes with washing buffer (0.3% Triton X-100 in PBS) before incubation with fluorescent secondary antibodies (Table 1) at room temperature for 1 hour. Finally, slides were washed 5 times for 5 minutes with washing buffer (0.3% Triton X-100 in PBS) before mounting with Fluoroshield Mounting Medium with DAPI (Abcam, Cat# ab104139) using 1.5 thickness coverslips. The following primary antibodies were used: Anti-Cd2ap (Proteintech, Cat# 51046-1-AP, RRID:AB_2879436); anti-TrkA (R&D, Cat# AF1056, RRID:AB_2283049); anti-choline acetyltransferase (ChAT; Sigma-Aldrich, Cat# AB144P, RRID:AB_2079751); anti-green fluorescence protein (GFP; Aves Labs, Cat# GFP- 1020, RRID:AB_10000240); and anti-Rab5 (Cell Signaling Technology, Cat# 46449, RRID:AB_2799303).

**Table 1.**
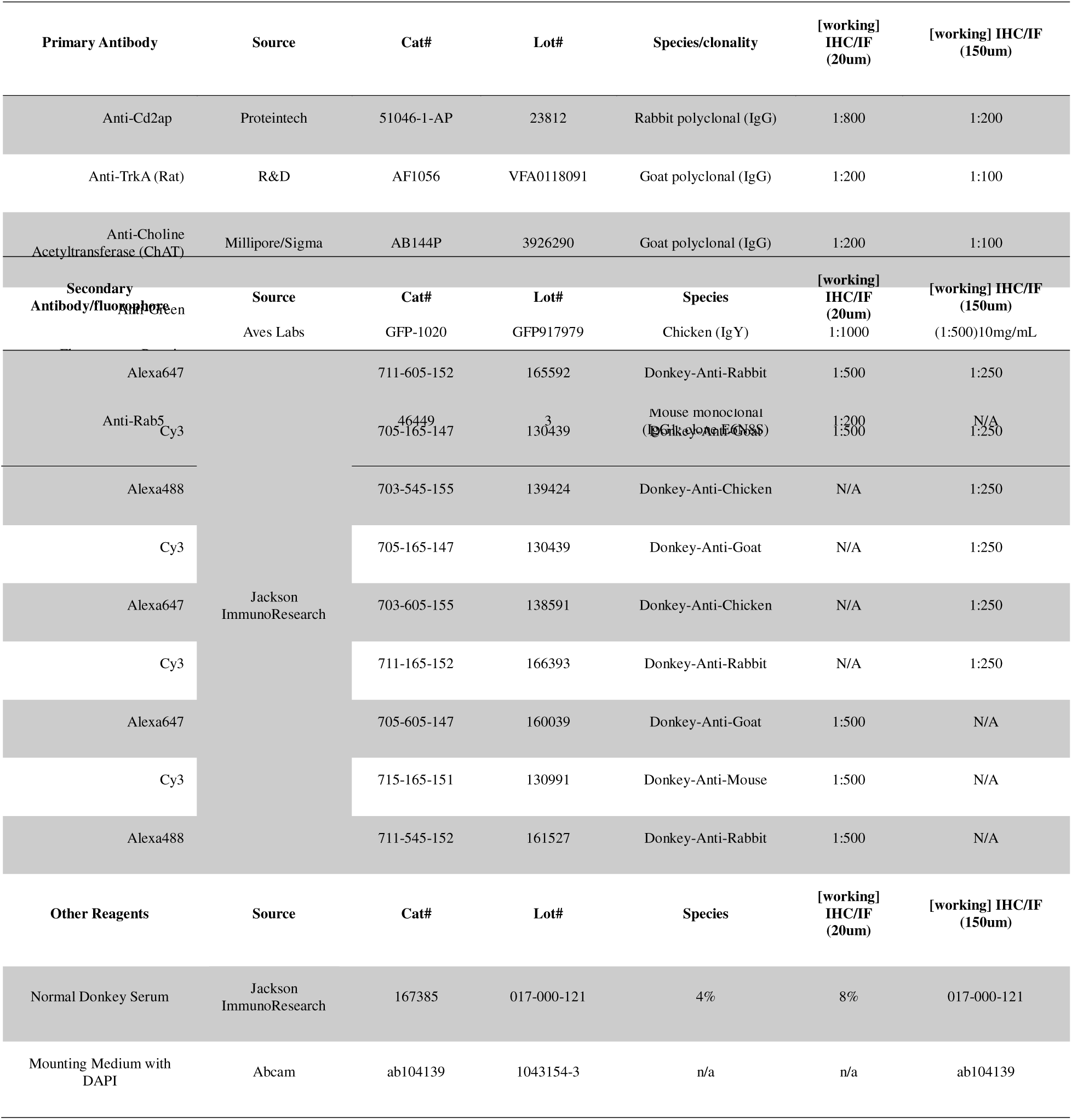
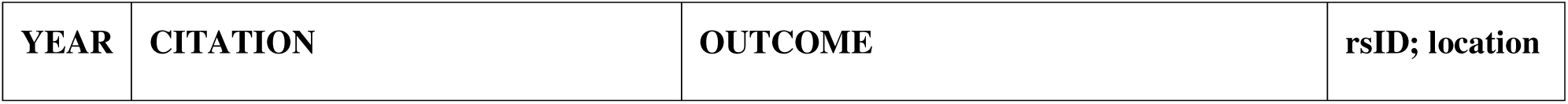
Antibodies and reagents used for immunofluorescence analysis.

### 2.5 Thick tissue Immunofluorescence (iDISCO)

Thick tissue sections (150µm) of ChAT-GFP mouse brains were embedded in 5% agarose in 1X PBS and cleared using iDISCO ^46^. Whole-section images of Chat-GFP+ brains were then imaged at 20X magnification using an epifluorescence microscopy slide scanner (BZ-X800 Slide Scanning Microscope, Keyence, USA). Concentrations of primary and secondary antibodies were increased for thick tissue section IF and are indicated in Table 1.

### 2.6 CD2AP Antibody Validation

The anti-Cd2ap primary antibody (Proteintech, Cat# 51046-1-AP, RRID:AB_2879436) was validated for use with qualitative and quantitative IF in two previous studies. Characterization of this antibody was performed using fixed, frozen sections of mouse kidney tissue, demonstrating signal specificity localized along the glomerular basement membrane, as expected ^47^. Subsequent validation of anti-Cd2ap was performed using Lentiviral-mediated, *Cd2ap*-knockdown tissue in human telomerase-immortalized retinal pigmented epithelial cells (hTERT-RPE1) as well as murine NIH3T3 cells ^48^. Investigators demonstrated that expression of CD2AP was localized to the primary cilium in control cells, and was ablated in knockdown tissues using both IF (See Figure 5, panels A-C, their manuscript) and Western Blot (See Figure 5, panel E of their manuscript) ^48^. Antibody specificity was confirmed in our laboratory using mouse kidney tissue (data not shown).

### 2.7 Quantification of CD2AP Fluorescence

#### Microscopy and Image Analysis

One slide/mouse (with all 15 sections) was imaged using an epifluorescence microscope (Leica DMi8, Leica Microsystems) at 20X. Four color micrographs from alternate (odd numbered) sections were then processed using FIJI software (ImageJ2; version 2.14.0/1.5f) ^49^. For fluorescence measurements in ChAT-GFP positive neurons, a binary mask was created by thresholding the GFP channel. The mask was processed to remove noise by removing foci<20μm in diameter. Masks were then layered over the ChAT antibody stain and then the CD2AP antibody stain to acquire the respective fluorescent signals from GFP-positive neurons.

#### Estimation of Background Fluorescence

Populations of cell bodies of developing cholinergic neurons undergo a phenotypic switch into adulthood, where multiple cholinergic cell subtypes are thought to be established based on synaptic properties ^50^. In ChAT-GFP+ mice, GFP signals are sustained into adulthood ^51^, and represent a subpopulation of cortical interneurons which synapse locally within the cortex, and are best characterized by their expression of vasoactive intestinal peptide ^43,50^. In the present study, we confirm that this population of adult cortical neurons remain ChAT-GFP positive, but do not stain with ChAT antibody. Therefore, this served as a measure of the background fluorescence.

### 2.8 Immunofluorescence Colocalization Assays

Colocalization assays were performed on IF z-stack, max intensity projection images collected using confocal microscopy (Leica Stellaris) and Image J *EzColocalization* plugin ^52^. To restrict analysis to Chat-GFP+ neurons, the polygon selection tool was used to trace the outline of the ChAT-GFP+ cell body from the GFP+ maximum projection, and the ‘region of interest’ (ROI) manager was employed to demarcate the GFP+ border onto the Cd2ap (yellow) and Rab5 (magenta) split channel maximal projections. Selection of images was restricted to those collected in the vDB region of the basal forebrain, and an average of 10 images per section, 20 sections per animal, in a total of 4 ChAT-GFP+ mice.

### 2.9 Statistical Analysis

All statistical analyses were conducted using Graphpad Prism 9. Data are presented as either individual animal (regional) mean values ± standard error of the mean (SEM) or the mean of all animals analyzed ± SEM. Data normality for individual animals as well as grouped data ware analyzed using the Kolmogorov-Smirnov test for data normalization, and was considered to be normally distributed when p >0.100. Protein expression, as reflected by fluorescence intensity was quantified as integrated density using FIJI (ImajeJ2). Mean CD2AP fluorescence intensity between brain regions was compared using a standard one-way analysis of variance (ANOVA) followed by Tukey’s post-hoc analysis. For immunofluorescence colocalization assays, degree of overlap for CD2AP and Rab5 colocalization was assessed using Pearson correlation analysis and the Costas coefficient and was restricted using FIJI to manually trace the soma of Chat-GFP+ cells and establish as a region of interest (ROI).

## Results

### Analysis of Publicly Available Single-Cell Sequencing Data Reveals that *Cd2ap* mRNA is Enriched in TrkA-Expressing Cholinergic Neurons

Our previous work provided one of the first descriptions of the structural and mechanistic dynamics of CD2AP in neurotrophin-mediated neuroplasticity, coordination of scaffolding interactions between TrkA and AKT subunit p85 in neurons and the positive regulation of retrograde signaling implemented through Rab5-decorated signalosomes ^5^. We further postulated that the retrograde, Rab5-mediated signaling mechanism, combined with the dynamics of the CD2AP-TrkA complex in cholinergic neurons would be a reasonable, potential mechanism(s), highly applicable to the pathogenesis of neurodegenerative disease like AD ^5^. Since this publication, germline variants in human *CD2AP* have remained at the forefront of GWAS investigating risk(s) for development of AD in humans ^16,53,54^.

The functional consequences of *Cd2ap* variants as well as the possibility of a potential role of CD2AP in CNS neurons is currently unknown. Using human AD GWAS studies, we first performed a review of the literature generated, with an emphasis on recent GWAS studies published between 2021 to present (date of submission, June 2024). Since 2021, we report a total of five GWAS studies for AD that identify *CD2AP* as a primary risk factor (and risk factor loci) for the development of AD in humans (Table 2). Of the loci identified, specific variant rs9349407 was correlated with increased risk for AD in Chinese populations ^55^ and specific variant rs7738720 was correlated with increased risk of AD pathophysiology in individuals of African ancestry ^15^. Overall, we identified a total of 19 GWAS studies performed worldwide (since the year 2010) where *CD2AP* loci have been positively associated with AD pathophysiology (Table 2).

**Table 2.** Summary of GWAS studies from 2013 to present (June 2024) implicating *CD2AP* in the pathogenesis and/or susceptibility to AD in humans.

|  |  |  |  |
| --- | --- | --- | --- |
| 2023 | Sherva R, Zhang R, Sahelijo N, et al. African ancestry GWAS of dementia in a large military cohort identifies significant risk loci. *Mol Psychiatry*. 2023;28(3):1293-1302. | Novel, significant SNPs near known AD risk genes TREM2, CD2AP, and ABCA7; peak variants in these genes differed from those previously reported in European and African ancestry cohorts. | rs7738720 |
| 2023 | Charisis S, Lin H, Ray R, et al. Obesity impacts the expression of Alzheimer's disease-related genes: The Framingham Heart Study. *Alzheimers Dement*. 2023;19(8):3496-3505. | Gene expression was measured for 74 AD-related genes, derived by integrating genome-wide association study results with functional genomics data from the Framingham Heart Study. Obesity metrics were associated with the expression of 21 AD-related genes. The strongest associations were observed with CLU, CD2AP, KLC3, and FCER1G. |  |
| 2022 | Gao S, Hao JW, Zhao YN, et al. An updated analysis of the association between CD2-associated protein gene rs9349407 polymorphism and Alzheimer's disease in Chinese population. *Front Neuroinform*. 2022;16:1006164. | Comparison of 11 independent studies from 8 articles exploring the correlation between rs9349407 variation and AD in Chinese. Additional meta-analysis based on fixed and random effect models were compiled to perform heterogeneity test. Use of additive model, dominant model, and recessive model for subgroup analysis demonstrated that the CD2AP rs9349407 polymorphism increases AD susceptibility in Chinese populations. | rs9349407 |
| 2022 | Khaire AS, Wimberly CE, Semmes EC, Hurst JH, Walsh KM. An integrated genome and phenome-wide association study approach to understanding Alzheimer's disease predisposition. *Neurobiol Aging*. 2022;118:117-123. | "Phenome-wide association" of 23 known, LOAD-associated SNPs and 4:1 matched control SNPs using UK Biobank data; associations confirmed known LOAD risk loci near PICALM, CD2AP, SPI1, and NDUFAF6, and identified a novel risk locus in epidermal growth factor receptor. |  |
| 2022 | Bellenguez, C., Küçükali, F., Jansen, I.E. et al. New insights into the genetic etiology of Alzheimer's disease and related dementias. *Nat Genet* 2022; 54:412–436 | Two-stage GWAS of 111,326 clinically diagnosed/'proxy' AD cases and 677,663 controls; found 75 risk loci, including 42 novel sites at the time of analysis. Pathway enrichment analyses confirmed involvement of amyloid/tau pathways and highlighted microglia implication. | rs7767350<br>6:47517390:T:C |
| 2021 | Li Y, Laws SM, Miles LA, et al. Genomics of Alzheimer's disease implicates the innate and adaptive immune systems. *Cell Mol Life Sci*. 2021;78(23):7397-7426. | Comprehensive review of GWAS studies (with meta-analyses from 2013 up to 2021) |  |
| 2021 | Kunkle BW, Schmidt M, Klein HU, et al. Novel Alzheimer Disease Risk Loci and Pathways in African American Individuals Using the African Genome Resources Panel: A Meta-analysis. *JAMA Neurol*. 2021;78(1):102-113. | Of 25 known AD loci in non-Hispanic, White individuals, only APOE, ABCA7, TREM2, BIN1, CD2AP, FERMT2, and WWOX were significantly associated with AD in African American individuals. |  |
| 2021 | Schwartzentruber J, Cooper S, Liu JZ, Barrio-Hernandez I, Bello E, Kumasaka N et al. Genome-wide meta-analysis, fine-mapping and integrative prioritization implicate new Alzheimer's disease risk genes. *Nat Genet* 2021;53(3):392–402. | AD GWAS with focus on causal genes and variants; confirmed recently discovered AD genes (CD2AP) also the top gene within 500 kb by network score and AD Risk. | rs9381563<br>6:47432637:C:T |
| 2019 | Jansen IE, Savage JE, Watanabe K, Bryois J, Williams DM, Steinberg S et al. Genome-wide meta-analysis identifies new loci and functional pathways influencing Alzheimer's disease risk. *Nat Genet* 2019;51(3):404–413. | AD GWAS of clinically diagnosed AD and AD-by-proxy cases; meta-analysis results revealed 29 risk loci, implicating 215 potential causative genes (CD2AP); analyses support a role for lipid processing and degradation of amyloid precursor proteins in AD pathogenesis. | rs9381563<br>6:47432637:C:T |
| 2019 | Kunkle BW, Grenier-Boley B, Sims R, Bis JC, Damotte V, Naj AC et al. Genetic meta-analysis of diagnosed Alzheimer's disease identifies new risk loci and implicates Abeta, tau, immunity and lipid processing. Nat Genet 2019;51(3):414–430. | AD GWAS; confirmed eQTLs for genes in previously identified loci, including *CD2AP* and the *CD2AP* locus | rs9473117<br>6:47431284:A:C |
| 2018 | Marioni, R.E., Harris, S.E., Zhang, Q. et al. GWAS on family history of Alzheimer's disease. Transl Psychiatry 2018; 8, 99. | Self-report of parental history of Alzheimer's dementia for GWAS of 314,278 participants from UK Biobank; meta-analysis with published consortium data revealed three new loci, relevant for AD and neurodegeneration, including *CD2AP*. | rs9381563<br>6:47432637:C:T |
| 2013 | Lambert JC, Ibrahim-Verbaas CA, Harold D, Naj AC, Sims R, Bellenguez C et al. Meta-analysis of 74,046 individuals identifies 11 new susceptibility loci for Alzheimer's disease. Nat Genet 2013; 45(12):1452–1458. | Meta-analysis of 17,008 cases and 37,154 controls, followed by replication in an independent cohort; confirmed previous GWAS findings (*CD2AP*) and identified 11 novel AD susceptibility loci: *HLA-DRB5-DRB1*, *SORL1*, *PTK2B*, *SLC24A4*, *ZCWPW1*, *CELF1*, *NME8*, *FERMT2*, *CASS4*, *INPP5D*, and *MEF2C* | rs10948363<br>6:47487762:A:G |

Although there is a clear association between *Cd2ap* variants and an increased risk of AD, the expression of *Cd2ap* (mRNA) or CD2AP (protein) in neurons adult CNS neurons in either mouse or human is poorly characterized. In order to determine and characterize CD2AP expression in CNS neurons, we performed an analysis of single-cell, neuronal, RNA-sequencing data publicly-available and sourced from DropViz webtool ^45^ (Figure 1A). Our initial analysis of neuronal cell types ordered neurons by level of CD2AP expression, expressed as transcripts per 100,000 in cluster. Of the few neuronal populations expressing CD2AP, these neurons also expressed the highest levels of *Ntrk1 (TrkA),* in cholinergic interneurons of the striatum, along with neurons located in the globus pallidus (GP). Striatum interneurons and GP neurons were found to also co-express the highest values of *Cd2ap* in the top-10, highest Ntrk1-expressing (cholinergic) neurons of the adult mouse brain. We next integrated a single cell RNA sequencing data from the Allen brain atlas (Figure 1 B)^56^. This dataset confirmed that CD2AP expression is most highly expressed in *Ntrk1*-expressing cholinergic neurons of the dorsal striatum and ventral striatum. Therefore, of all neuronal cell types present in the adult mouse brain, *Cd2ap* mRNA was enriched in TrkA-expressing, cholinergic neurons.

**Figure 1.**
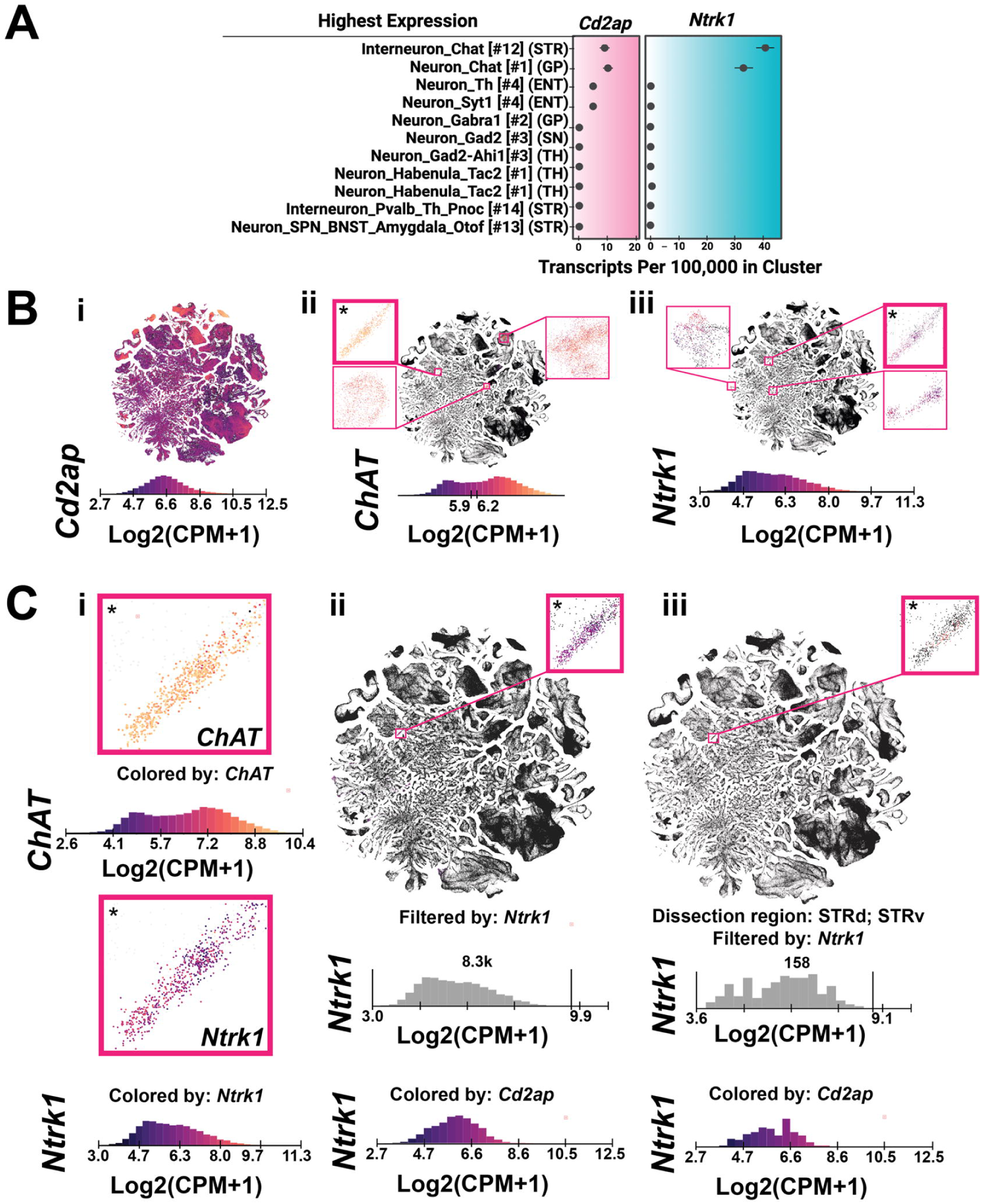
Neuronal CD2AP mRNA is enriched in *Ntrk1* (TRKA)-expressing, cholinergic neurons of the adult mouse brain. (A) Summary of single-cell RNA-sequencing data from the adult mouse brain, using the Dropviz webtool (Saunders *et al,* 2018). The top ten, highest *Cd2ap*-expressing neuron populations are shown on the left (magenta) and corresponding expression levels of *Ntrk1* on the right (cyan), indicates that CD2AP is enriched in *Ntrk1*-expressing cholinergic neurons. (B) Whole brain scRNAseq data from the Allen Mouse Brain Atlas^56^, clustered into neuronal subtype and shaded according to expression of *Cd2ap* (B_i_), *ChAT* (B_ii_), and *Ntrk1* (B_iii_). * Clusters with increased expression in *Chat*^+^/Ntrk1^+^/*Cd2ap*^+^ neurons in dorsal striatum (STRd) and striatum ventral striatum (STRv). (C_i_) Comparison of *ChAT* and *Ntrk1* expression profiles from the STRv/STRd. (C_ii_) Co-expression of *Ntrk1* with *Cd2ap.* (C_iii_) Co-expression of *Ntrk1* with *Cd2ap*.

### CD2AP Protein is Expressed in Cholinergic Neurons of the Murine Basal Forebrain

To verify that CD2AP protein is expressed in ChAT neurons, (as predicted by analysis of online gene expression data, in Figure 1), we performed an immunofluorescence analysis of frozen tissue sections from ChAT-GFP reporter mouse brains (Figure 2). Whole-section analysis of ChAT-GFP brains confirmed the bilateral presence and expression of ChAT-GFP positive neurons throughout the superior thalamic radiation (striatum), nucleus accumbens and substantia innominata (Figure 2, a).

**Figure 2.**
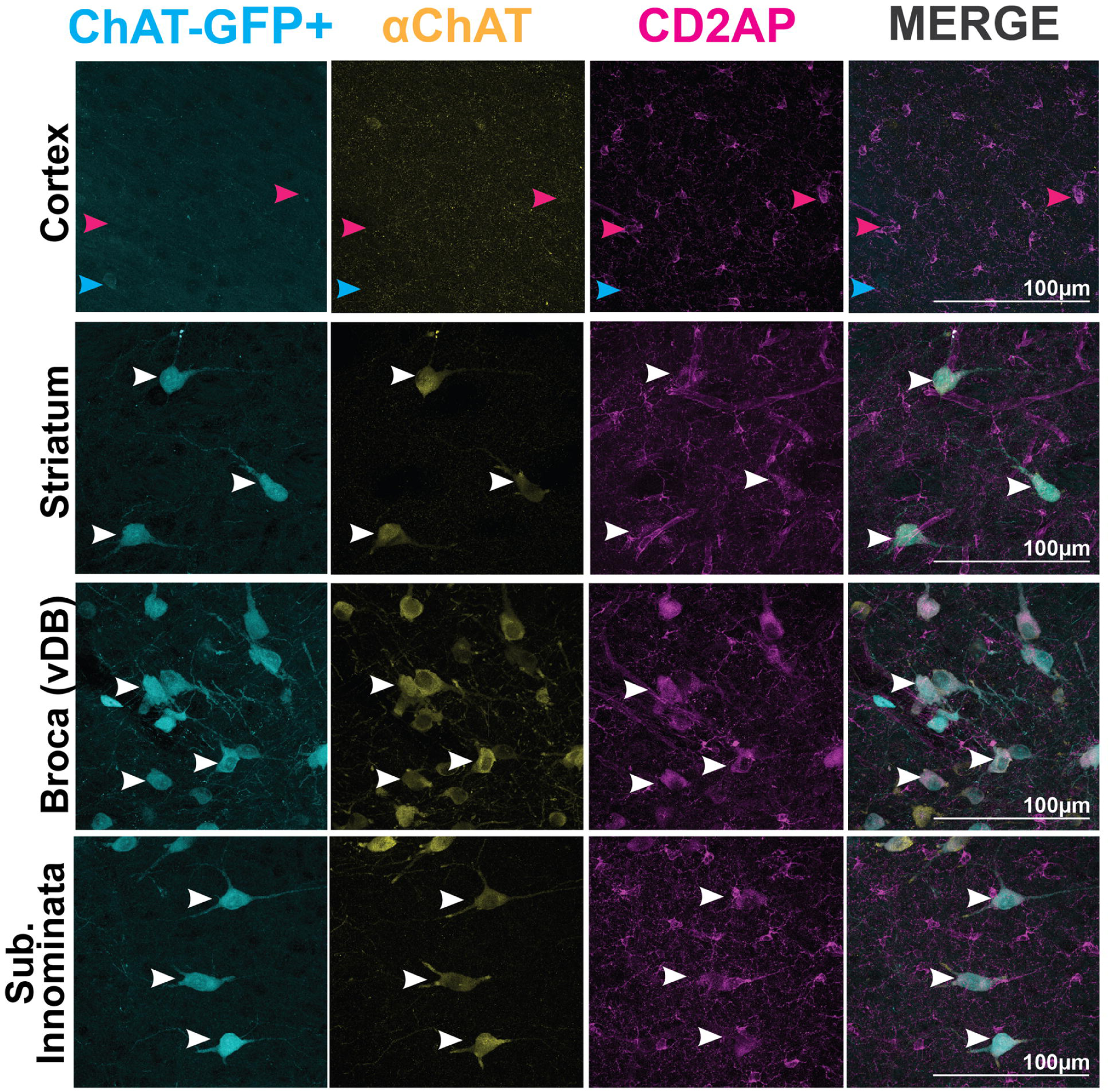
Immunofluorescence analysis of CD2AP protein expression in ChAT-GFP neurons in regions of interest within the adult mouse brain. Representative immunofluorescence confocal stacks from 20um-thick frozen tissue sections. ChAT-GFP signal (left column, cyan) was detected in ChAT-lineage neurons (ChAT-GFP+/11ChAT-) in Cortex (top row), Striatum, Broca (vDB) and Substantia Innominata. Fluorescence signal from ChAT-antibody (11ChAT) (second column, yellow) was observed in neuron soma of adult (ChAT-GFP+/ 11ChAT+) cholinergic neurons in Striatum (second row), Broca (third row) and Substantia Innominata (bottom row). Fluorescence signal of CD2AP (third column, magenta), was observed in adult cholinergic neurons (ChAT-GFP+/ 11ChAT+), but not in cholinergic-lineage neurons (ChAT-GFP+/11ChAT-) in the cortex. Cyan arrows indicate ChAT-GFP+/11ChAT-/CD2AP- cholinergic lineage neuron cell bodies; magenta arrows indicate (cortical) vascular 11CD2AP staining; white arrows indicate ChAT- GFP+/11ChAT+/CD2AP+ neuron cell bodies.

CD2AP expression was highest in the basal forebrain region, where we observed the presence of ChAT-GFP neurons (cyan) throughout the vDB. Co-localization of ChAT-GFP with anti-ChAT antibody (Figure 2, yellow) in cholinergic neurons of the vDB confirmed the specificity of the ChAT primary antibody. Note, absence of ChAT antibody staining in Chat-GFP+ cell bodies specifically in the cortical region also confirmed the specificity of the anti-ChAT antibody. As documented by previous characterization of the ChAT-GFP transgenic mouse brain, developmental expression of GFP reporter persists in the adult^57^. ChAT is expressed during a developmental window in cortical neurons, and therefore, in the adult these neurons contain ChAT-GFP reporter but conversely are ChAT-antibody negative ^57^. Therefore, this confirms the specificity of the CD2AP antisum and served as a convenient proxy to quantify background fluorescence.

Next, to quantify expression of CD2AP in cholinergic populations, confocal scans were collected from the following regions of adult mouse brains: Substantia innominata (basal forebrain) region, vDB, the superior thalamic radiation (striatum), and the cortex. Three-channel fluorescence signals were acquired from ChAT-GFP reporter, CD2AP antibody and ChAT antibody (Figure 3). Double-positive signal from CD2AP and ChAT antibodies, were detected in the majority of ChAT-GFP+ cell bodies in the basal forebrain, vDB and striatum. ChAT-GFP+ soma in the cortex region did not stain with the anti-ChAT antibody (yellow), again validating the specificity of immunofluorescent staining. Note that although CD2AP signal was not present in the cell bodies of cortical CHAT-GFP neurons, it was expressed in cortical vascular structures and non-neuronal cells, as expected^58^, providing additional evidence of antibody specificity. Taken together, these results validate the specificity of CD2AP immunostaining, and qualitatively show that CD2AP is co-expressed with ChAT in the substantia innominata (basal forebrain), vDB, and striatum.

**Figure 3.**
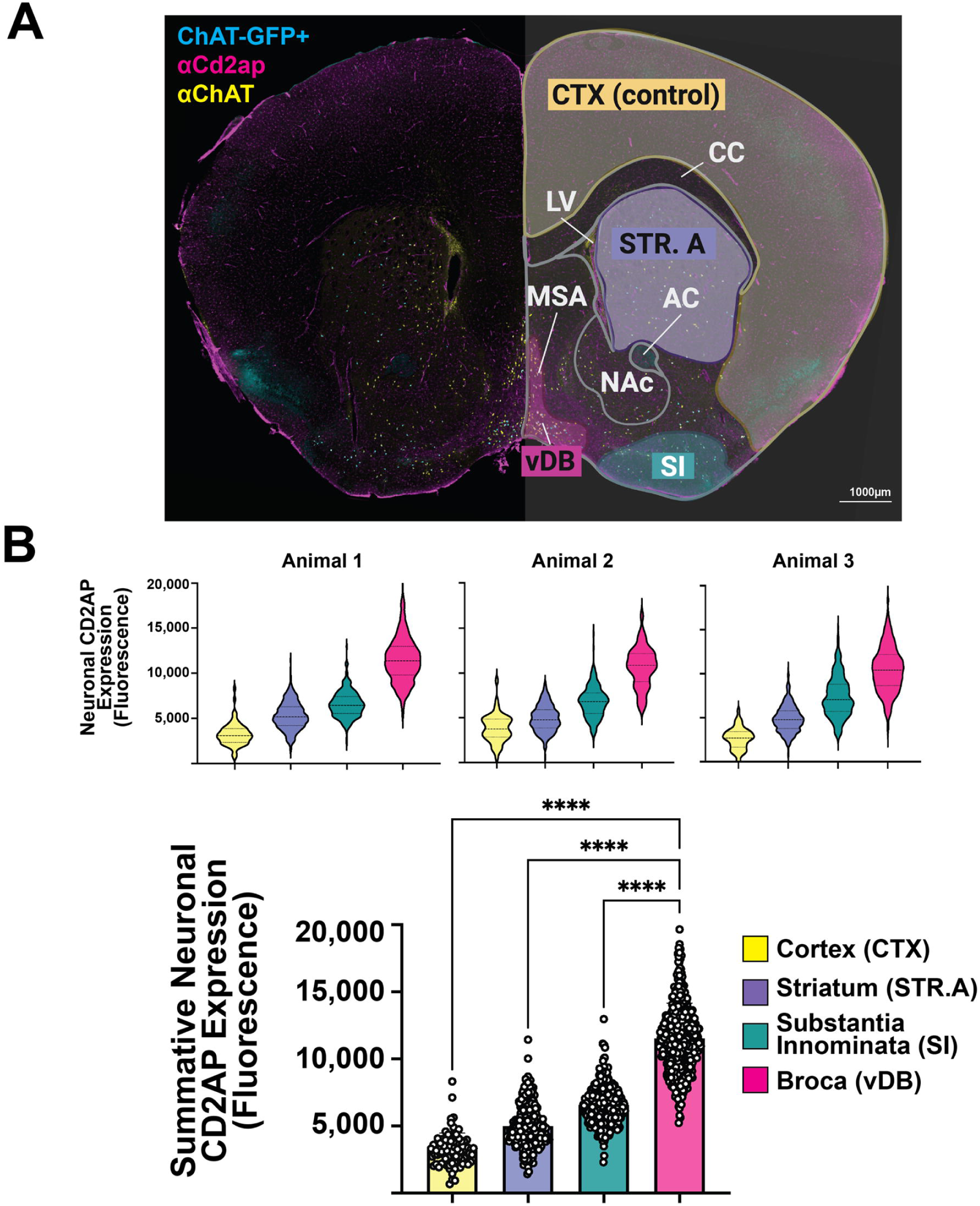
Quantification of CD2AP protein immunofluorescent signals in cholinergic neurons. (A, left) Representative coronal tissue section (20um) of ChAT-GFP mouse brain stained with primary antibodies to ChAT (yellow) and Cd2ap (magenta). (A, right) Regions of interest to be quantified are indicated and labeled, where regional color key corresponds to the quantification grafts quantified in Figure 3B. (B, top) Mean CD2AP immunofluorescence signals in ChAT-GFP+ soma from indicated brain regions, collated from 3 animals. (B, bottom) Mean, regional expression levels (expressed as integrated density) of Cd2ap in individual mice; violin plots clusters reflect raw data point showing within-animal distribution and mean values ± standard deviation. One-way analysis of variance (ANOVA) with Tukey post-hoc analysis revealed Cd2ap expression in the basal, striatal and optic regions of the mouse brain, with cortex serving as a background-signal control. ****p<0.0001.

**Figure 4.**
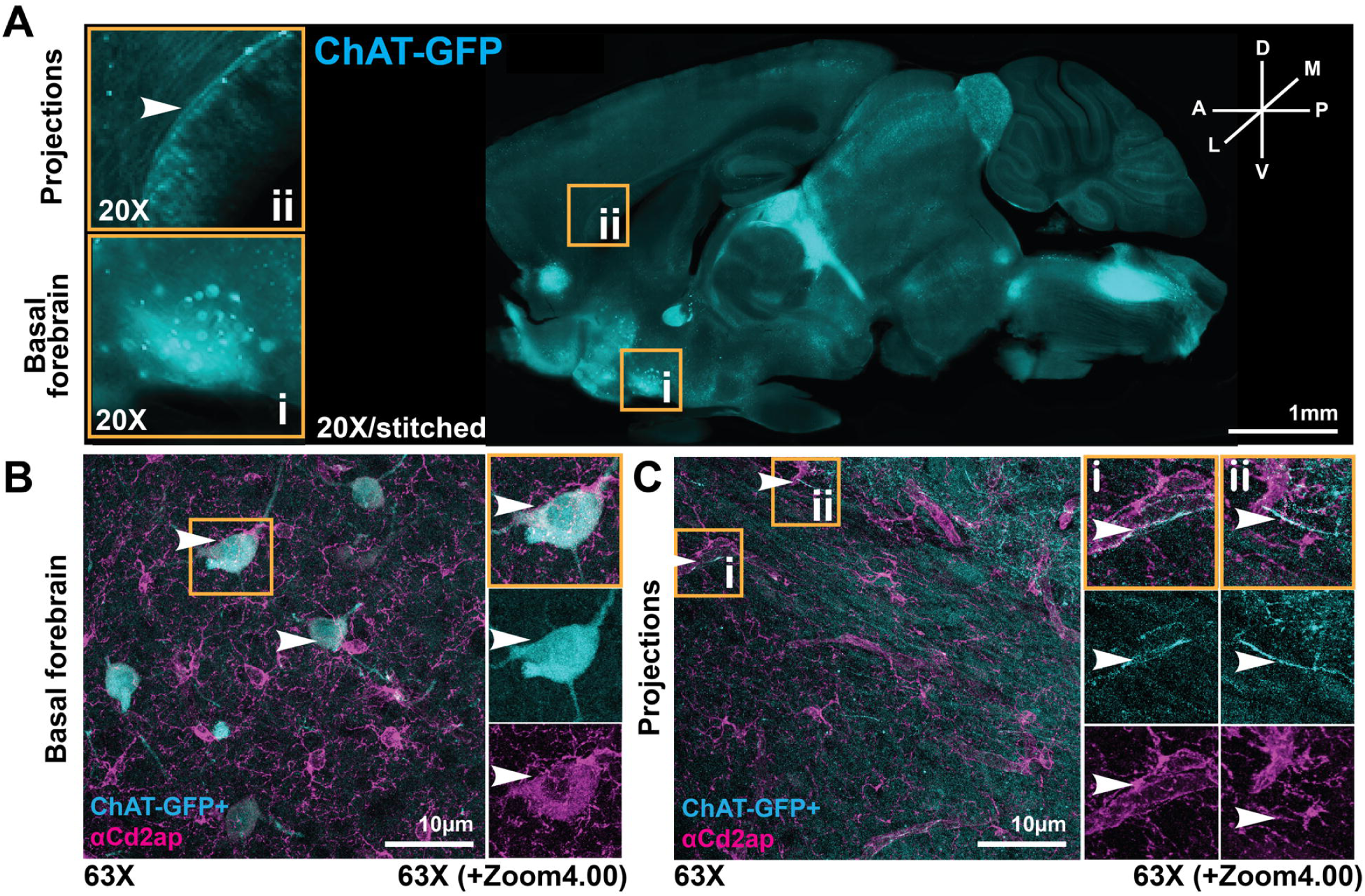
CD2AP expression in ChAT-GFP+ adult mouse brain projections. (A) Representative, whole-brain thick slice imaging of ChAT-GFP reporter, sagittal sections (150um) embedded in agarose. (A; i) ChAT-GFP+ cholinergic neuron cell bodies originating in the basal forebrain; (A; ii) ChAT-GFP+ axonal projections (white arrow) from basal forebrain on trajectory towards hippocampal/iso-cortical regions. (B-C) Representative, regional examination of CD2AP in confocal slices of ChAT-GFP+ neuron cell bodies in the basal forebrain (B; white arrows) shows staining patterning consistent with ChAT-GFP+ axonal projections (C; i, ii). Yellow box indicates inset; cyan is ChAT-GFP+ fluorescence signal; magenta is signal from anti-Cd2ap immunofluorescence; white arrows indicate areas of overlap between ChAT-GFP and Cd2ap.

Next we quantified CD2AP fluorescence intensity in the cortex (negative control), superior thalamic radiation, substantia innominata (basal forebrain) and Broca’s area (vDB) (Figure 3B). CD2AP fluorescence (integrated density) values of individual animals were expressed as mean values ± SD, and collated group regional data were expressed as mean values ± SEM. In the grouped regional mean data, the lowest CD2AP fluorescence (mean integrated density ± SEM) was observed in the cortex (3449 ± 124), serving as a baseline/background measurement. Mean signal intensity from the superior thalamic radiation was 4936± 27.6, the substantia innominata (basal forebrain) was 7428 ± 55.9 and Broca’s area/vDB was 10929 ± 72.3. A one-way ANOVA with Tukey post-hoc analysis revealed a statistically significant difference among brain regions (p=0.0004). CD2AP fluorescence signals were significantly higher in the Broca/vDB regions when compared to background (cortex) (p=0.0085). Mean CD2AP expression in the vDB was also statistically significantly greater when compared to the superior thalamic radiation (p=0.0018), and the basal forebrain (p=0.04).

### CD2AP is Locates to Cortical Projections from Basal Forebrain Neurons

We observed an overlap in signaling between CD2AP and ChAT-GFP extending from cell bodies in the basal forebrain continuing throughout the neuronal projections extending anteriorly and superiorly in proximity to the hippocampus and within the isocortex (Figure 5). As previously reported, vascular expression of CD2AP is substantial^23^ and was clearly visualized as such throughout all regions of the mouse brain. However, CD2AP co-labeling in ChAT-GFP projections was observed in a smaller/finer distribution, clearly distinct from CD2AP expression in blood vessels (Figure 5, bottom).

**Figure 5.**
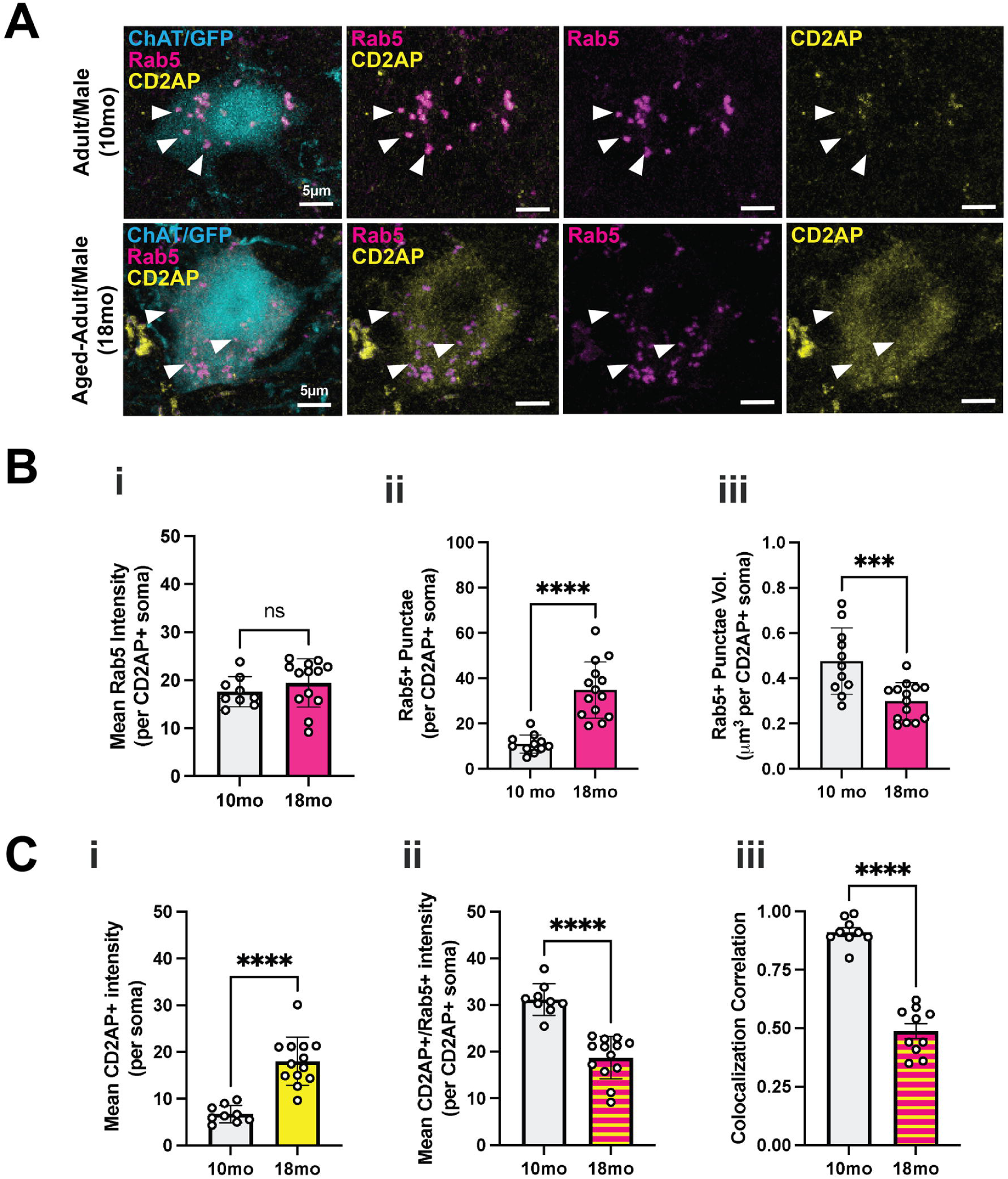
Cd2ap/Rab5 co-expression and morphology in aged male mice. (A) Representative maximum intensity projections of Cd2ap (yellow) and Rab5 (magenta) in ChAT-GFP reporter cholinergic neurons (cyan) of the VDB in adult males (10 mo; top row) and in aged-adult male mice (18 mo; bottom row). White arrowheads indicate representative CD2AP+/Rab5+ puncta of ChAT- GFP+ soma. (B) Quantitative analysis of (i) mean Rab5 expression (expressed as fluorescence intensity per CD2AP+ soma), (ii) number of Rab5+ puncta (per CD2AP+soma), and (iii) volume of Rab5+ puncta (expressed as µm^3^ per CD2AP+ soma) in 10 and 18 month-old male mice. (C) Quantitative analysis of (i) CD2AP expression (expressed as fluorescence intensity per soma), (ii) mean coexpression of CD2AP+/Rab5+ expression (expressed as fluorescence intensity per CD2AP+ soma), and (iii) Pearson correlation analysis of CD2AP+/Rab5+ colocalization in 10 and 18 month- old male mice. Not statistically significant (ns); p<0.0001; p<0.001.

### CD2AP is colocalized with Rab5 Endosomes in Cholinergic Basal Forebrain Neurons

The Rab-GTPase Rab-5 is required for NGF-TrkA retrograde trophic signaling in sensory neurons^59^ and in TrkA-expressing neurons of the CNS that are primarily cholinergic basal forebrain neurons^60^. Morphological abnormalities in early endosomes are an early pathological marker in dementia, where dysfunctional rab5-decorated vesicles appear enlarged and accumulate in large quantities within the cell soma^61,62^. Misregulated Rab5 is associated with cholinergic neuron degeneration and memory impairment, where Rab5 puncta increase in size and accumulate in cholinergic soma (e.g,. ^63^).

Upregulation of basal forebrain rab5 expression also correlated with Braak staging in prodromal AD^64^, and choline supplementation ameliorates early endosome pathology in basal forebrain cholinergic neurons in mouse models of Down syndrome and AD^65^. Therefore, targeting mechanisms of RAB5 protein in endo-lysosomal impairment is an emerging therapeutic goal (e.g., ^33^).

Confocal imaging revealed punctate CD2AP staining morphology in cholinergic neurons (Supplemental Figure 1, Figure 5). Our previous studies in NGF-responsive sensory neurons, demonstrated that CD2AP is an adaptor protein that coordinates retrograde endosome signaling via interaction with rab-GTPases^5^. Therefore, we characterized and quantified CD2AP colocalization with Rab5, a small GTPase, mediator and sorter of endosomal formation in cholinergic neurons. Using confocal microscopy, analysis of immunofluorescence signals from ChAT-GFP reporter mice revealed an abundance of CD2AP and Rab5 colocalization in punctae, present in ChAT-GFP+ neurons in the basal forebrain tissue (Figure 5A). In order to quantify colocalization of CD2AP with Rab5, we developed a colocalization assay in ChAT-GFP cell bodies, revealing a near-complete overlap between CD2AP and Rab5 punctuates with mean Pearson correlation R values of 0.80 and Costas P value of 1.00 in adult mice (Supplemental Figure 1). In light of our previous studies in sensory neurons, these observations suggest that CD2AP-adaptor protein may be coordinator of retrograde NGF-TrkA signaling in cholinergic neurons.

### 3.5 CD2AP expression is increased and Rab5 colocalization decreased in ChAT-GFP+ Basal Forebrain Neurons in a cohort of aged male mice

It is established that endocytosis proteins accumulate during ageing in the normal human brain and increases in endocytosis may be a potential mechanism of neuronal aging^66^. RAB5 puncta size and number are increased in cholinergic basal forebrain neurons of aging mice (e.g., ^67^) and in middle-aged Human brains prior to changes in Aβ levels^68^. We performed analyses of RAB5 morphology in a cohort of adult mice (10 months old; N=6) compared to aged-adult (18 months old; N=6) male mice (Figure 5). In accordance with previous studies, we observed that the number of Rab5+ puncta increased significantly from 10 to 18 months (p<0.0001; Figure 5Bii) in cholinergic neurons and the mean volume of Rab5+ puncta decreased significantly with age (p<0.001; Figure 5Biii), whereas Rab5 fluorescence intensity (expressed as mean fluorescence intensity in CD2AP+ soma) did not significantly change with age (Figure 5Bi).

Qualitative examination of Chat-GFP+ soma in the vDB of adult and aged-adult male mice revealed apparent differences in the endosomal numbers and general morphology of Rab5+/CD2AP+ puncta per soma (Figure 5A). CD2AP expression appeared to become more diffuse and less punctate, in addition to a decreased in degree of overlap (colocalization) of CD2AP with Rab5 aged adults when compared to adult male mice (Figure 5A). When quantifying expression of CD2AP, and in contrast to Rab5 expression, we found that mean CD2AP intensity (expressed as fluorescence intensity, per soma) increased significantly from 10 to 18 months (p<0.0001; Figure 5Ci). We found that coexpression of CD2AP with Rab5 decreased with age (p<0.0001; Figure 5Cii), and colocalization correlation also decreased (p<0.0001; Figure 5Ciii). These initial observations suggest that the functional interaction between CD2AP adaptor scaffold and Rab5 GTPase may become dystrophic with age in cholinergic neurons, and may therefore point to a mechanism by which CD2AP gene variants may contribute to risk of dementia.

## Discussion

By analysis of publicly available scRNA-seq data from the central nervous system, we observed elevated expression of *Cd2ap* mRNA in *Ntrk1*-expressing (NGF dependent) cholinergic neurons. Using an *in vivo* reporter mouse for cholinergic neurons, we identified and quantified the expression of CD2AP protein with ChAT-GFP reporter and ChAT-antibody staining in cholinergic neurons of the superior thalamic radiation (striatum), nucleus accumbens/substantia innominata, the basal forebrain, and the vDB, most pronounced of which was quantified in the vDB/basal forebrain.

Expression of CD2AP in ChAT-GFP projections from the basal forebrain to the (iso-) cortex, suggested a role for CD2AP in retrograde axonal transport signaling, and for maintaining basal forebrain communication with higher regions. Co-staining with Rab5 revealed near-complete colocalization between CD2AP and Rab5 punctuate patterning, consistent with a role for CD2AP in NGF-TRKA retrograde signaling, as previously characterized in primary sensory neurons and in cholinergic neurons *in vitro* ^5,36^. In this study, we are the first to characterize the expression of the Alzheimer risk locus, *Cd2ap,* in populations of adult CNS neurons *in vivo*. Additionally, we propose that association between CD2AP and Rab5 in these neurons is likely disordered in aging mouse brains, suggesting a role for CD2AP in retrograde trophic signaling in cholinergic neurons.

CD2AP was first characterized as an adhesion molecule critical for the structural integrity, filtration and overall function of the kidney, as determined using various genetic knockout/knock-down studies *in vivo* and *in vitro* ^22,28,69^. Interestingly, reduced kidney function and diagnosis with chronic kidney diseases (CKD) is an established risk factor for cognitive decline and AD, although the mechanism has yet to be fully elucidated ^70–72^. The biochemistry and function of CD2AP was first characterized in T-lymphocytes of the immune system, where it was described as an adhesion molecule responsible for mechanisms of protein segregation and cytoskeletal polarity, and where it was first postulated to orchestrate dynamics of receptor patterning via the cytoskeletal scaffold ^1^. This initial characterization prompted further exploration of CD2AP expression. In other cell and organ systems, and specifically in the podocyte cells of the kidney, a highly-specialized, gate-keeping cell found in the glomerulus ^1,69–72^. In podocytes, CD2AP is located to the actin-rich scaffolds of the slit-diaphragm, a charge-selective opening through which vascular/metabolic waste products are filtered from circulating blood and excreted from the body in urine. Loss of CD2AP is homozygous-lethal, due to the rapid-onset of severe nephrotic syndrome and glomerular disease, resulting in kidney failure and death in mice by 6 weeks of age due to failure of the slit-diaphragm ^22,69^. Thus, CD2AP functions as a mediator and organizer of F-Actin structures, where it colocalizes with Rab4 in podocyte endosomes ^73,74^. Genetic mutations and/or impairment of CD2AP, therefore, not only impairs cellular cytoarchitectural scaffold, but also decreases endosomal-mediated degradation processing, influencing secretase and amyloid precursor proteins (APP) transport and ultimately enabling the accumulation of APP segments and tangles ^33,75,76^.

CD2AP is also required for the maintenance of structure and permeability of systemic blood vessels (e.g. epithelial and endothelial cells) ^22,58,77^. Interestingly, analysis of microvessels isolated from human patients with confirmed AD observed Cd2ap expression to be linearly associated with cognitive decline and Cd2ap KO from brain microvasculature (microvessels) in mice resulted in cognitive impairment ^58^. Interaction between the vasculature and the immune system has also been implicated in the onset, progression and severity of AD ^78,79^. In vascular endothelial cells CD2AP regulates intercellular adhesion molecule-1 (ICAM-1) ^27^. Loss of CD2AP initiates increased clustering of ICAM-1 on the endothelial cell surface, leading to excessive leukocyte transmigration and subsequent pro-inflammatory signaling pathways associated with kidney dysfunction, arthritis and atherosclerosis ^27^. CD2AP has also been shown to be highly expressed in plasmacytoid dendritic cells (pDC), a specialized subtype of dendritic cells known to initiate type-1 interferon signaling in response to inflammation and in microbial pattern recognition ^80^.

In addition to its roles in renal, vascular and immune cells, CD2AP was identified as a genetic risk factor for the development of AD in humans ^81,82^, a finding that has been supported by a substantial number of studies across model organism species, including *Mus musculus* ^58^, *Drosophila* ^32,41^, and *Caenorhabditis elegans* ^83,84^. Previously established *Cd2ap* variants have even been incorporated along with variants in *Trem2^em1Adiuj^, App^em1Adiuj^* and *Apoe^tm^*^1^.^1^*^(APOE*4)Adiuj^* into a multi-genetic/causal, humanized mouse model of AD (JAX Strain# 033873) ^85,86^. Although CD2AP is known to be expressed in tissues and cell types outside of the kidney (e.g. primary sensory neurons ^5^, endo/epithelial cells ^22,23^ and migratory glial cells ^58,87^), inherent expression of CD2AP in cholinergic neurons of CNS (*in vivo*) has not previously been identified or quantified, supporting the plausibility of a pathogenic mechanism of AD inherent to cholinergic neurons of the CNS.

Early characterizations of CD2AP (previously also referred to as ‘Cas ligand with multiple Src homology (SH) 3 domains’ (CMS)) identified that coexpression of active Rab4 and CD2AP *in vivo* resulted in the enlargement of early endosome formations, was required (along with c-Cbl) for proper endosomal formation, and finally, that CD2AP interactions with both Rab4 and Rab7 might also be involved in degradation of these endosomes ^38^. Interestingly, CD2AP expression in conditionally immortalized mouse podocytes was shown to be directly coexpressed with Rab4, but not Rab5, in late-endosomal processing ^74^. However, subsequent research from our laboratory and others, has indicated that CD2AP interaction with specific Rab isoforms may, in fact, vary by cell/tissue type, where NGF signaling via TrkA receptor in primary, DRG neurons, is dependent upon TrkA+/Rab5+ endosomes for mediation of long-range NGF signaling ^5,37^. In the present study, we document the visual and quantitated colocalization of CD2AP and Rab5 *in vivo,* where previous research has identified interactions with Rab3 ^48^, Rab4 ^37^, and RIN ^2,36^ or isolated with Rab5 *in vitro* ^2,37^. Evidence in support of a role for Rab5 specifically for endosomal formation and the propagation of trophic signaling in cholinergic neurons ^5,33,88^, in conjunction an established association of impaired Rab5 signaling in brain tissue of deceased humans with confirmed AD ^35^, are consistent with the findings of the present study, and further support the hypothesis that CD2AP-Rab5 endosomal signaling in cholinergic neurons of the basal forebrain is a viable, mechanistic explanation for AD pathogenesis.

In the nervous system, CD2AP plays a regulatory role in neuronal cell survival mechanisms, a process that is known to be trophic-factor dependent ^6^. Early investigations into the possible pathological relevance of trophic signaling in AD showed that while retrograde transport was reduced, there is a specific reduction in NGF retrograde transport observed from the hippocampus to the septum in basal forebrain cholinergic neurons in a Ts65Dn mouse model of Down’s Syndrome, also known to be applicable to mouse models of AD ^89^. In addition to what is known about CD2AP, Rab5-mediated endocytosis is also hypothesized to be dysregulated in AD ^35,63^, a finding that is also supported by human GWAS studies ^12,90^.

## Conclusion

Disruption of endocytosis in cholinergic neurons is key to the pathophysiology of AD, and may lead to accumulation of amyloid beta and/or the formation of neurofibrillary tangles. In this study we highlight and summarize GWAS studies supporting the idea that variants in genes known to be critical for endocytosis and endocytic processes (like *Cd2ap* and *Rab5*) are consistently associated with increased susceptibility to AD. We demonstrated that CD2AP is expressed in both the cell body and projections of basal forebrain cholinergic neurons in the CNS, where it collocates with RAB5 GTPase-decorated signalosomes, and we provide histological evidence suggesting that CD2AP- RAB5 interaction may become disordered in ageing animals. Together, these data support our working hypothesis that CD2AP may orchestrate endocytosis in cholinergic neurons and may therefore play a role in healthy aging and/or the pathogenesis of dementias such as AD.

CD2AP function has been characterised in multiple cell types critical for the function of various tissues/organs like those of the kidney, vasculature, immune system, sensory neurons of the PNS. and now, cholinergic neurons of the CNS. Taken together, we propose a working model through which CD2AP-coordinated signaling may serve as a viable target to sustain endocytosis and trophic signaling in neurons, and as a multiorgan/wholistic approach to improving AD outcomes.

## Author Contributions

All authors were involved in experimental design and analysis and manuscript preparation.

## Acknowledgments

Animal care services were provided by UNE COBRE Behavior Core; histology & imaging training and services were provided by the UNE COBRE Histology & Imaging Core Facilities. Both UNE Behavior Core and Imaging & Histology Core facilities are also supported by the UNE Center for Pain Research (NIGMS UNE COBRE #P30GM145497; PI: Ian Meng). The content is solely the responsibility of the authors and does not necessarily represent the official views of the National Institutes of Health. We wish to thank Christoph Straub for providing numerous tissues and experimental assistance as well as members of the Harrison lab who assisted with imaging, experiments and laboratory management, especially Peter Neufeld, BS.

## Data Availability Statement

The data that support the findings of this study are available from the corresponding author upon reasonable request.

