## Supplementary figures and images for "Adult Neuronal Expression of the Alzheimer Risk Protein CD2AP is Enriched in NGF-Responsive Basal Forebrain Neurons, where it Collocates with Rab-5 Endosomes"

### supplimental method

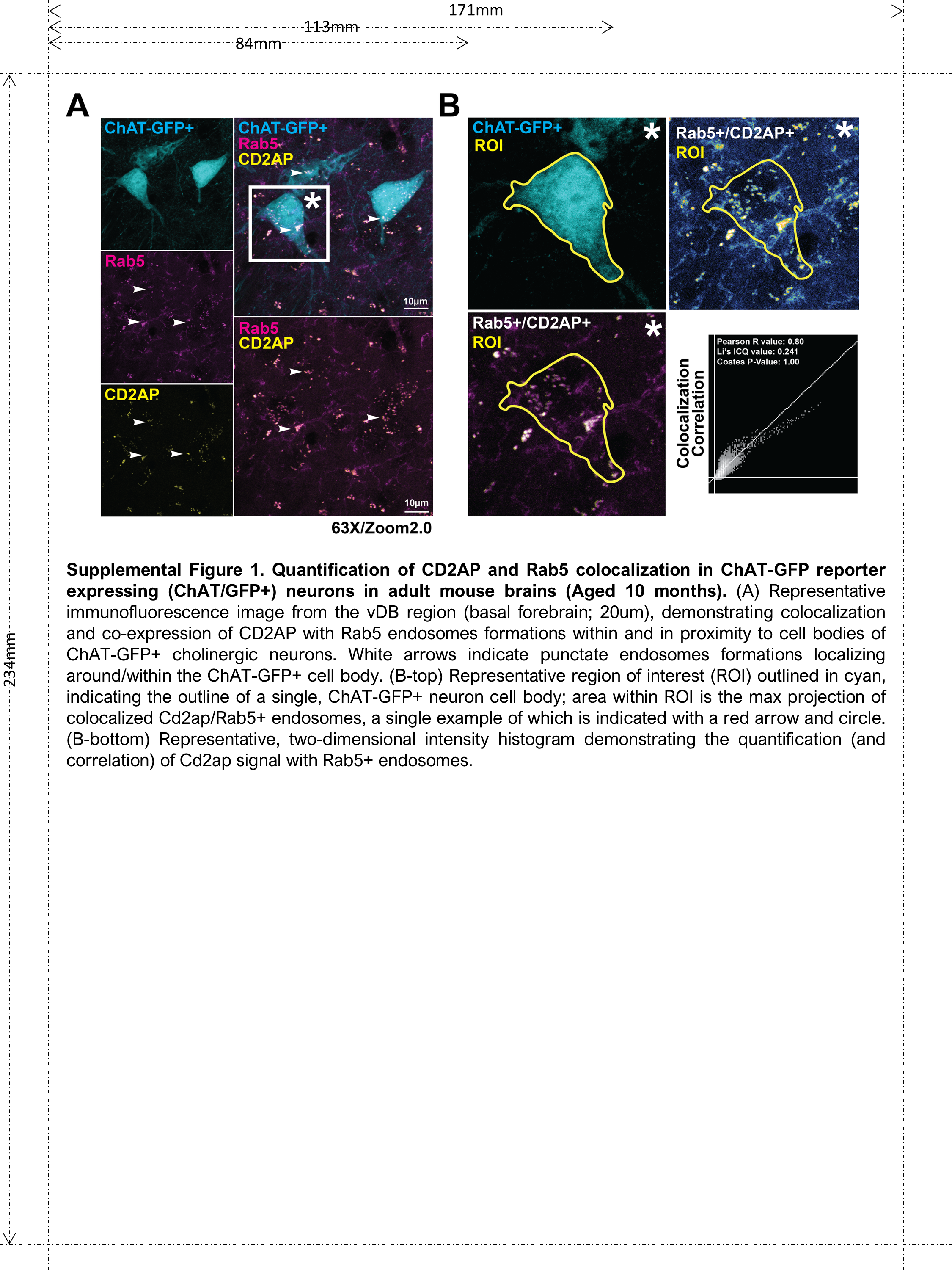
